# Vacuum Insulated Probe Heated ElectroSpray Ionization source (VIP-HESI) enhances micro flow rate chromatography signals in the Bruker timsTOF mass spectrometer

**DOI:** 10.1101/2023.02.15.528699

**Authors:** Mukul K. Midha, Charu Kapil, Michal Maes, David H. Baxter, Seamus R. Morrone, Timothy J. Prokop, Robert L. Moritz

**Affiliations:** Institute for Systems Biology

**Author notes:** Address correspondence to Robert Moritz.

**Keywords:** Vacuum Insulated Probe Heated ElectroSpray Ionization source, CaptiveSpray, Electrospray ionization, Micro-flow rate Liquid chromatography, Data Independent Acquisition- Parallel Accumulation–Serial Fragmentation, Slice-Parallel Accumulation–Serial Fragmentation

## Abstract

By far the largest contribution to ion detectability in liquid chromatography-driven mass spectrometry-based proteomics is the efficient generation of peptide ions by the electrospray source. To maximize the transfer of peptides from liquid to a gaseous phase to allow molecular ions to enter the mass spectrometer at micro-spray flow rates, an efficient electrospray process is required. Here we describe superior performance of new Vacuum-Insulated-Probe-Heated-ElectroSpray-Ionization source (VIP-HESI) coupled with micro-spray flow rate chromatography and Bruker timsTOF PRO mass spectrometer. VIP-HESI significantly improves chromatography signals in comparison to nano-spray ionization using the CaptiveSpray source and provides increased protein detection with higher quantitative precision, enhancing reproducibility of sample injection amounts. Protein quantitation of human K562 lymphoblast samples displayed excellent chromatographic retention time reproducibility (<10% coefficient-of-variation (CV)) with no signal degradation over extended periods of time, and a mouse plasma proteome analysis identified 12% more plasma protein groups allowing large-scale analysis to proceed with confidence (1,267 proteins at 0.4% CV). We show that Slice-PASEF mode with VIP-HESI setup is sensitive in identifying low amounts of peptide without losing quantitative precision. We demonstrate that VIP-HESI coupled with micro-flow-rate chromatography achieves higher depth of coverage and run-to-run reproducibility for a broad range of proteomic applications.

## Introduction

Large scale proteomic analysis for biomarker studies, as one example, requires consistent interrogation of assembled cohort samples in a highly reproducible and high-throughput fashion to obtain quantitative differences with strong statistical significance of perturbed networks of proteins. Nano-flow chromatography-tandem mass spectrometry (nLC-MS/MS), a mainstay of proteomics analysis, provides high-sensitivity with minimal sample usage, but its quantitative proteomic application is restricted by poor robustness over extended periods of time leading to low sample throughput and additional batch effect considerations^1^. As next-generation mass spectrometer (MS) instruments become increasingly faster and more sensitive, microflow chromatography (μLC) workflows circumvent such issues by attaining stable electrospray ionization, higher signal stability, resulting in higher data reproducibility with the ease of use as compared to nanospray ionization^2–8^. Consequently, microflow chromatography-tandem mass spectrometry (μLC–MS/MS) has gained an advantage over nLC-MS/MS in large-scale proteome research experiments where sample amounts are not limited and have shown to achieve a significant proteome coverage^7^. The highly reproducible peptide measurements by μLC–MS/MS is further improved with the advancements in the data-independent acquisition (DIA) methods that can achieve high protein depth and coverage, with excellent reproducibility compatible with μLC–MS/MS^9^.

Different high-flow rate chromatography setups have been used with a variety of mass spectrometer instruments. For instance, we and several others have applied capillary flow rates coupled with TripleTOF mass spectrometers (SCIEX), accelerating data measurements for increased sample throughput^3,10–15^. Lately, several studies have shown the application of microflow rate^1,16^, and capillary flow^9^ chromatography workflows in robustly quantifying thousands of proteomes using a set of Thermo-Fisher Orbitrap mass spectrometers. More recently, the Bruker timsTOF MS system, with added ion mobility, has gained popularity due to its higher duty cycle and Parallel Acquisition and Serial Fragmentation (PASEF) mode which provides an additional dimension of peptide ion separation that results in a reduction in the sample complexity and achieves deep proteome coverage^17^. Szyrwiel *et al*. benchmarked the Slice-PASEF method using analytical flow (500 μL/min) coupled with a new Vacuum Insulated Probe Heated ElectroSpray Ionization (VIP-HESI) source and timsTOF MS^18^; however, to achieve the balance between performance and robustness, μLC–MS/MS could be more suitable for proteomics applications^7^. This led us to evaluate the performance of microflow chromatography for quantitative proteome analysis in Slice-PASEF and dia-PASEF modes using a sensitive timsTOF ion mobility MS. For most proteomics applications, the Bruker timsTOF MS is coupled with nanoflow rate chromatography through the CaptiveSpray (CS) ion source. To achieve more robust operation and ease of use, we investigated the performance of the new VIP-HESI ion source from Bruker as well as the standard ElectroSpray ionization (ESI) operated in μLC–MS/MS mode and compared it with a CS ion source using nanoflow rate workflows.

In our comparative analysis, we standardized the VIP-HESI source with microflow chromatography workflow with two different analytical column dimensions (*i.e*., 0.5 and 1.0 mm ID) operating at 20 μL/min and 40 μL/min flow rates, to investigate the efficiency of the proteomic analysis of synthetic peptide mixtures (PepCalMix), HeLa (an immortalized human epithelial cell line) and K562 cell line tryptic digests, and tryptic digests of undepleted mouse plasma using dia-PASEF. The acquired data illustrates higher sensitivity and higher reproducibility in peptide ion MS and MS/MS collection using VIP-HESI than compared to standard ESI and CS setups. These results suggest that the VIP-HESI source with microflow chromatography using DIA methods in timsTOF MS has the potential of processing thousands of samples in an automated unattended operation with superior higher performance characteristics.

## Experimental Procedures

### Preparation of PepCalMix synthetic peptide mixture

An isotopically light version of the Sciex PepCalMix solution containing 20 non-naturally occurring synthetic peptides that cover a wide mass and retention time range (SynPeptide, Shanghai, China). A stock solution of 1 pmol/μL was prepared in 5% v/v acetic acid in 2% v/v acetonitrile (ACN)-containing water and aliquots of 10 μL each was stored at −80° C until further use. For ESI and CS-MS measurements, 1 μL of the PepCalMix aliquot was diluted in 39 μL of the same solvent to achieve the final peptide concentration: 25 fmol/μL. For VIP-HESI MS linearity measurements, 10 μL of the PepCalMix aliquot was diluted in 790 μL in the same solvent to achieve a final concentration of 12.5 fmol/μL.

### Mouse tissue and plasma sample preparation

#### Mouse husbandry and interventions

All the mouse samples were prepared at the University of Michigan in a specific pathogen-free colony kindly supplied by Dr. R. Miller. Genetic background and husbandry conditions for the Snell dwarf and GHRKO mice were described previously^19^. Mice used for Rapamycin (encapsulated, used at 14 ppm), Canagliflozin (180 ppm), 17α-estradiol (17aE2) (14.4 ppm), Acarbose (1000 ppm), and Calorie-Restricted samples were of the UM-HET3 stock, produced as the offspring of CByB6F1/J mothers and C3D2F1/J fathers, as described in this article^20^. The base diet was Purina 5LG6. To monitor specific-pathogen status, sentinel mice were exposed to spent bedding for two weeks prior to testing and all tests were negative for the entire aging colony during the experimental period. The protocols were reviewed and approved by the University of Michigan’s Institutional Animal Care and Use Committee. The metadata of the mouse liver, kidney, gastrocnemius muscle tissues, and plasma samples are provided in **Supplementary Table 1.**

#### Sample Harvesting

Mice were humanely euthanized at 12 months of age. Euthanasia occurred between 8 am and 11 am (lights cycled on at 6 am and off at 6 pm) by rapid asphyxiation in a bag containing CO_2_ gas. Mice were unconscious within 5 seconds and stopped breathing at about 10 seconds. The interval between removal from the home cage to death was less than one minute. Mice were removed from the bag as soon as breathing ceased, and blood was harvested by closed-chest cardiac puncture.

#### Sample Preparation

Tissue sections from kidney, liver, and gastrocnemius muscle were processed for DIA LC-MS/MS analysis as follows. Frozen tissue was lysed in 50 mM ammonium bicarbonate buffer with 5% sodium dodecyl sulfate (SDS) and homogenized on a Precellys Evolution (Bertin Instruments, France) for 9 rounds of 20 sec at 8500 rpm with 30-sec rest in between. Protein concentration of the lysate was determined using the BCA protein assay (Pierce, Cat# 23227). Lysate containing 200 μg protein was aliquoted out and denatured for 2 min at 90° C. After denaturation, samples were reduced with 5 mM TCEP ((tris(2-carboxyethyl)phosphine) Sigma Cat# 4706) for 30 min at 60° C and then alkylated with 10 mM ɑ-iodoacetamide (Millipore Cat# 407710) for 30 min at room temperature in the dark. Samples were acidified with phosphoric acid to a final concentration of 1.2% and S-Trap buffer (100 mM ammonium bicarbonate buffer + 90% methanol) was added at 1:7 ratio prior to loading on S-Trap wells (96-well plate format, ProtiFi, USA) according to the manufacturer’s instructions. Using a positive pressure manifold (Resolvex M10, Tecan, USA) samples were washed twice with S-Trap buffer before overnight digestion at 37° C with Trypsin (Promega Cat# V511X) in digestion buffer (50 mM ammonium carbonate) at 1:50 ratio. Digested peptides were eluted as per the manufacturer’s instructions (S-Trap 96-well plate protocol version 1.4). Importantly for normalization, all plates contained an even mix of samples from all tissues, arranged systematically to avoid bias across either rows or columns and in nearly all cases all three tissue samples from the same donor were loaded onto the same plate.

Mouse undepleted plasma from 284 mice was processed for DIA LC-MS/MS analysis as follows. Frozen plasma aliquots were thawed at 4° C for 3 hours followed by a hard spin to pellet insoluble particles (5 min, 10000 x g, 4° C). Protein concentration was determined using the bicinchoninic acid (BCA) protein assay (Pierce, Cat# 23227). Samples were normalized by aliquoting 400 μg in phosphate-buffered saline (PBS) into a final volume of 37.5 μL. Denaturation was performed by adding a 12.5 μL lysis buffer (200 mM Triethylammonium bicarbonate (TEAB), 20% SDS) to a final concentration of 5% SDS and heating for 5 min at 90° C with 800 rpm shaking. All mixing and shaking steps were performed in a ThermoMixer C orbital thermal controlled shaker (Eppendorf) with the Smartblock for deep-well plates. After denaturation, samples were reduced with 5 mM TCEP (Sigma Cat# 4706) for 15 min at 55° C and then alkylated with 10 mM iodoacetamide (Millipore-Sigma Cat# 407710) for 10 min in the dark at room temperature, both with 800 rpm shaking. Samples were acidified with orthophosphoric acid to a final concentration of 2.7% (v/v). S-trap buffer (100 mM TEAB pH 8.5/90% methanol) was added at a 1:7 ratio prior to loading on S-trap 96 well-plate format (ProtiFi, USA) by 1 min spin at 4000 x g. Once loaded, samples were washed six times with 400 μL S-trap buffer and spun at 4000 x g for 5 min to ensure complete dryness.

Tryptic digestion was performed at 37° C with 125 μL Trypsin (Promega Cat# V511X) in digestion buffer (100 mM TEAB pH 8.5) at a 1:25 ratio. After the first hour of incubation at 37° C, an additional 75 μL digestion buffer was added to prevent the drying out of the samples. After overnight incubation at 37° C, the digested peptides were eluted with 80 μL digestion buffer and centrifuge at 4000 x g for 1 min, and then with 80 μL 50% ACN in digestion buffer and centrifuge at 4000 x g for 1 min. The final eluate was ~250 μL. Quantification of the digested peptides was performed by the Fluorescamine fluorescent peptide assay^21^ (Pierce, Cat# 23290) before drying down to completion.

#### HeLa cell culture and protein digestion

HeLa S3 immortalized human epithelial cells (ATCC Cat No. CCL-2.2) were cultured in Dulbecco’s Modified Eagle’s Medium supplemented with 10% fetal bovine serum (FBS) were grown at 37° C in 5% CO_2_ to confluency. Cells were rinsed with cold PBS three times, frozen on ethanol/dry ice in pellets, and stored at −80° C. Cells were lysed in urea buffer (8 M urea, 0.1 M Tris pH 8.2) by three rounds of flash-freezing in ethanol/dry ice followed by thawing/vortexing. The lysate was sonicated on ice for 8 rounds of the 30 s in a cup-horn sonicator (full power) to shear any DNA, then centrifuged for 15 min at 21,000 x g. Protein quantity was determined by BCA assay. The lysate was reduced with 5 mM TCEP for 30 min at 37° C in a benchtop incubator at 850 rpm, alkylated with 15 mM iodoacetamide for 30 min at 20° C in the dark at 850 rpm, and then quenched with 5 mM TCEP for 30 min at 20° C at 850 rpm. Protein was precipitated in 80% ethanol and Automated Protein-Aggregation Capture (PAC) was performed on a BioSprint-96 (Qiagen) according to the following method: plate 1 - magnetic head comb, plate 2 - 1.6 μg/μL carboxylate beads (1:1 mixture of hydrophilic and hydrophobic beads [mass/mass], GE Healthcare Cat. Nos. 24152105050250 and 65152105050250) in 500 μL H_2_O per well, plate 3 - 500 μL 80% ethanol per well, plate 4 - 0.4 μg/μL lysate in 500 μL 80% ethanol per well, plate 5 to 7 - 500 μL 80% ethanol per well, plate 8 - 300 μL 50 mM ammonium bicarbonate per well. The program was set to collect the beads from plate 2, wash for 5 min at medium speed in plate 3, bind protein for 15 min at medium speed in plate 4, wash for 5 min per plate at medium speed for plates 5 through 7, and mix beads for 10 min at medium speed in plate 8. The beads were then left in plate 8, transferred to 1.5 mL microcentrifuge tubes, and trypsin (Promega) at a ratio of 1:100 (w/w) trypsin: protein lysate was added to the tubes. Digestion was carried out for 18 hours at 37° C on a tube rotator. Digestion was quenched with the addition of formic acid to 1%, beads were resolved on a magnet, and the supernatant was dried in a SpeedVac (Thermo-Fisher). Samples were resuspended in 0.1% formic acid\H_2_O (v/v) for liquid chromatography-mass spectrometry.

#### K562 cell culture and protein digestion

K562 cells (ATCC CCL-243, human bone marrow myeloid leukemia lymphoblast cell line) were cultured at 37 °C with 5% CO_2_ in Eagle’s minimum essential medium ((EMEM), ATCC 30-2003) supplemented with 10% FBS and were grown to 70% confluence. Cells were lysed in urea buffer (8 M urea, 0.1% RapiGest, and 100 mM ammonium bicarbonate pH 8). Protein quantity was determined by BCA assay. Proteins were reduced with 5mM TCEP for 30 min at 37° C in a benchtop incubator at 850 rpm, alkylated with 10 mM iodoacetamide for 30 min at room temperature in the dark, and digested using trypsin (1:50 w/w), and samples desalted with tC18 SepPak cartridges (Waters).

#### High-pH Fractionation

To maximize the proteome coverage from HeLa digest, peptides were fractionated using a reversed-phase Acquity CSH C18 1.7 μm 1 × 150 mm column (Waters, Milford, MA) on UltiMate 3000 high-pressure liquid chromatography (HPLC) system (Dionex, Sunnyvale, CA) operating at 30 μL/min and SpotOn (Bruker). 5 mM ammonium bicarbonate as buffer A and 100% ACN as buffer B were used. Peptides were separated by a linear gradient from 5% B to 35% B in 55 min, followed by a linear increase to 70% B in 8 min. 24 in-solution fractions were collected in a concatenated manner^22^, acidified to pH < 2 with trifluoroacetic acid, and vacuum dried prior to DDA (data dependent acquisition) LC-MS/MS.

#### Liquid chromatography-mass spectrometry

All peptide samples were spiked in with iRT standard peptides (Biognosys AG, Schlieren, Switzerland) and applied to MS analysis using EasyLC nano HPLC (Thermo-Fisher Scientific), Vanquish Neo HPLC system (Thermo-Fisher Scientific), Evosep One system HPLC (EvoSep) and nano Elute HPLC (Bruker) configured in either nanoflow or microflow modes were coupled to a timsTOF PRO mass spectrometer (Bruker). Both modes were operated with 99.9% water, 0.1% formic acid/Milli-Q water (v/v, buffer A), and 99.9% ACN, 0.1% formic acid (v/v, buffer B). Four different HPLC systems, four different samples, three different ion sources, nine different analytical columns, and eight different gradient lengths were used in this study **(Supplementary Table 2)**.

For CS measurements of PepCalMix and mouse plasma, nanoflow mode was selected, and peptides were trapped on a 0.5 cm × 0.3 mm trap cartridge Chrom XP C18, 3 μm (Thermo-Fisher Scientific) at 10 μL/min and separated on C18 UHP 15 cm × 0.15 mm × 1.5 μm column (Bruker/PepSep) at 1 μL/min using a 45-minute stepwise gradient from 3% to 25% B in 37 min, 25% to 35% B in 8 min, 35% to 80% B in 1 minute and isocratic flow at 80% B for 2 min followed by fast washing and equilibration for three column volumes. For generating a comprehensive HeLa sample library, peptides were separated on C18, 8 cm × 0.15 mm × 1.5 μm column (Bruker/PepSep), C18, 15 cm × 0.15 mm × 1.5 μm column (Bruker/PepSep), C18, 15 cm × 0.15 mm × 1.9 μm column (Bruker/PepSep), C18, 25 cm × 0.2 mm × 1.5 μm column (Bruker/PepSep), C18, 25 cm × 0.15 mm × 1.9 μm column (Bruker/PepSep), C18, 25cm × 0.075mm × 1.6μm column (IonOpticks, Australia), C18, 40cm × 0.075mm × 1.9 μm column (Bruker/PepSep) using premade 200SPD, 100SPD, 60SPD, 30SPD, and 15SPD gradient methods using Evosep One and 90- minute linear gradient from 3% to 35% B in 90 min, 35% to 95% B in 20 minute and isocratic flow at 95% B for 20 min followed by equilibration of column for starting conditions for 9 minutes. The CS ion source was equipped with a 20 μm emitter (Bruker) and the parameters were as follows: 1700 V Capillary voltage, 3.0 L/min dry gas, and temperature set to 180° C.

For ESI and VIP-HESI measurements, the flow path was changed to microflow in the Vanquish Neo, peptides were trapped on a 50 x 1 mm ID trap cartridge Chrom XP C18, 3 μm (Thermo-Fisher Scientific) at 50 μL/min and separated on either C18, 15 cm × 1 mm × 1.7 μm Kinetix column (Phenomenex, USA) at 40 μL/min using the same 45 minutes stepwise gradient as described above or C18 20 cm × 0.5 mm × 1.9 μm column (Dr. Maisch GmbH) at 20 μL/min using a linear 21-minute gradient from 3 to 35% B in 21 min, 35 to 80% B in 1 minute and isocratic flow at 80% B for 2 min. The standard ESI source was equipped with a 100 μm electrode probe (Bruker) and the parameters were as follows: 4500 V Capillary voltage, 10.0 L/min dry gas, and temperature 200° C. The VIP-HESI spray ion source was equipped with a 50 μm electrode probe (Bruker) and the parameters were as follows: 4000 V of Capillary voltage, 3.0 L/min Dry gas, and temperature 200° C, probe gas flow 3.0 L/min and temperature 100° C.

PepCalMix, mouse liver, kidney, gastrocnemius muscle tissues, and plasma sample injections were acquired using Bruker timsTOF preformed dia-PASEF mode schema covering the *m/z* range of 400-1200 and 1/K0 range 0.6 to 1.42 in 32 × 25 Da windows with a mass overlap of 1 Da, resulting in a total cycle time of 1.8 s. A total of seven different concentrations of PepCalMix peptides (12.5 fmol, 25 fmol, 50 fmol, 100 fmol, 200 fmol, 400 fmol, and 800 fmol), and six different injection amounts (0.4 μg, 2 μg, 4 μg, 10 μg, 20 μg, and 40 μg) of mouse plasma sample and 200 ng each of liver, kidney, gastrocnemius muscle tissues samples were measured. For the Slice-PASEF experiment, three different concentrations of HeLa injections *viz*. 10 ng, 100 ng, and 1000 ng were measured. The 1Frame (1F) Slice-PASEF method was downloaded from the recently published study^18^ that covers the 400 to 1000 *m/z* and the 1/K0 range 0.75 to 1.2. The premade py8 dia-PASEF method (that was compared with the 1F Slice-PASEF method) was modified to set the precursor mass range to 400 to 1000 *m/z* and 1/K0 0.65 to 1.37, with 24 × 25 Da windows with ramp and accumulation time 72 ms and total cycle time estimated to 0.7 s. For both methods, high sensitivity mode was enabled in the tims control acquisition software of the mass spectrometer.

For HeLa library generation, 24 fractions were measured using timsTOF MS (Bruker) set to dda-PASEF acquisition scan mode covering 100–1700 *m*/*z* with 10 PASEF ramps. The TIMS settings were 100 ms ramp and accumulation time (100% duty cycle), resulting in 1.1 s of total cycle time. Active exclusion was enabled with a 0.4 min release. The default collision energy with a base of 0.6 1/*K*0 [V s/cm2] is set at 20 eV and 1.6 1/*K*0 [V s/cm2] at 59 eV was used. Isolation widths were set at 2 *m*/*z* at <700 *m*/*z* and 3 *m*/*z* at >800 *m*/*z*. To achieve more comprehensive coverage, HeLa digest peptides were also acquired using dia-PASEF preformed py3, py5, and py8 (Bruker) schema based on the length of gradient used in the Evosep One HPLC.

#### Spectral assay library generation

For detailed quantitative proteomic analysis in individual experiments, we generated two comprehensive hybrid sample-specific spectral libraries for both HeLa and mouse plasma samples using DDA and DIA runs in Spectronaut^23^ 16.0.220606.53000 (Biognosys) respectively. The mouse spectral library was built using search archives (.psar) files from a published study^24^ and in-house sample-specific DIA acquisition files of mouse liver, kidney, gastrocnemius muscle tissues, and plasma samples. The HeLa library was generated using 24 fractionated DDA files from the HeLa sample and sample-specific DIA HeLa sample files as detailed in **Supplementary Table 2**. For respective libraries, related acquisition files were searched against the Mouse UniProt one protein sequence per gene FASTA (downloaded on March 07, 2021, containing 21,991 entries) and Human UniProt FASTA (downloaded on October 21, 2022, containing 20,375 entries). For each library iRT peptides (Biognosys) were appended and carbamidomethyl was used as fixed modification (C); acetyl (protein N-term), oxidation (M) as a variable modification; and enzyme digestion trypsin/P with up to two missed cleavages were allowed. Mass tolerances were automatically determined by Spectronaut, and for the rest of the processing, default settings were used. Identification search results were filtered at a 1% false discovery rate on the PSM, peptide, and protein levels. Sample-specific PepCalMix library was generated against the PepCalMix peptide FASTA sequence using the Spectronaut 17.0.221202.55965 (Biognosys) with the same setting described above.

#### Spectral assay library quality assessment using DIALib-QC

Both comprehensive spectral assay libraries were assessed for their quality using DIALib-QC (v1.2)^25^. DIALib-QC evaluates 57 parameters of compliance and provides a detailed report of the library’s complexity, characteristics, modifications, completeness, and correctness. The assessment reports are provided in **Supplementary Table 3**. In the DIALib-QC assessment report, there were no problem assays found for both libraries, and were used as it is for DIA-MS analysis.

#### Data Analysis

Two different DIA software tools were used in this study, Spectronaut (Biognosys, Switzerland) to process PepCalMix data & mouse plasma samples, and DIA-NN^26^ to perform the targeted data extraction of Slice-PASEF HeLa data files.

#### Spectronaut

Quantification and DIA processing of PepCalMix and control UMHET mouse plasma samples were performed using Spectronaut DIA software (version 16.0.220606.53000 (Biognosys, Switzerland). The mouse assay library was used directly as generated and described above. For the non-linear iRT calibration strategy, a dynamic window was used for both mass tolerance (MS1 and MS2), and to set up the extracted ion chromatogram (XIC) retention time (RT) window. Pre-processing of MS1 and MS2 calibration strategies was enabled. Decoy assays were dynamically generated using the scrambled decoy method with a set size of 0.1 as a fraction of the input library size. The identification was performed using the kernel density estimator with precursor and protein identification results filtered with a q-value of <0.01. For quantification, MS2 ion peak areas of quantified peptides were averaged to estimate the protein peak areas. Additional parameter settings were used as default.

For PepCalMix sample data processing, the PepCalMix library was directly used and the identification was performed using a normal distribution density estimator with precursor and protein identification results filtered with a q-value of <0.01. For quantification, the Top N ranking order setting was disabled to include all 20 peptides to estimate PepCalMix protein quantity. For the linearity experiment, all seven different concentrations were analyzed together and for direct comparison of PepCalMix protein quantities between different sources, joint processing of the respective measurements was performed. For both Spectronaut sessions, using grid view the estimated quantity profile data in log2 scale was exported.

Large-scale 284 plasma sample data analysis was performed using Spectronaut DIA software (version 17.0.221202.55965 (Biognosys, Switzerland) using both mouse library (as described above) and directDIA (library-free mode) to increase the proteome coverage and reduce the sparsity in the combined data matrix. For directDIA workflow, the database and parameter settings were kept the same as described above. Default settings were used without global normalization enabled. Trypsin specificity was set to two missed cleavages and a false discovery rate of 1% on both peptide and protein were used. Data filtering was set to q-value. For processing K562 samples, directDIA (library-free mode) was implemented within Spectronaut as described above.

#### DIA-NN

DIA-NN 1.8.2 beta 11 version^18^ was used for the targeted extraction of three replicates of each injection amount for both Slice-PASEF and dia-PASEF measurements. The comprehensive HeLa assay spectral library was provided to DIA-NN and library precursors were annotated with the human FASTA database. The mass accuracies were set to 15 ppm (both MS1 and MS2) and the scan window was fixed to 5. The protein inference was disabled and spectral library protein grouping information was used. For Slice-PASEF analysis, --tims -scan was provided to DIA-NN in the additional options section. The measurements corresponding to each combination of the acquisition method and the injection amount were analyzed separately, and the output .tsv file was filtered at a 1% global protein q-value.

## Results and Discussion

### Linearity assessment using VIP-HESI source and its comparative performance with standard ESI and CS ion sources using 20 PepCalMix peptides

The linear relationship between the signal response of the mass spectrometer and concentration using VIP-HESI ion source was studied. We collected seven different concentration points of a pool of 20 synthetic peptides (PepCalMix protein) to cover a wide range of 12.5 fmol to 800 fmol (see Experimental Procedures). An excellent linear gain in the total protein quantity was observed as we increased the PepCalMix concentration with the goodness of fit R-Squared (R^2^) of more than 99% **(Fig. 1a and Supplementary Table 4a))**. The linear response enabled fair comparison between the CS, ESI and VIP-HESI sources, and the response range used is shown with excellent fit to linearity.

**Figure 1.**
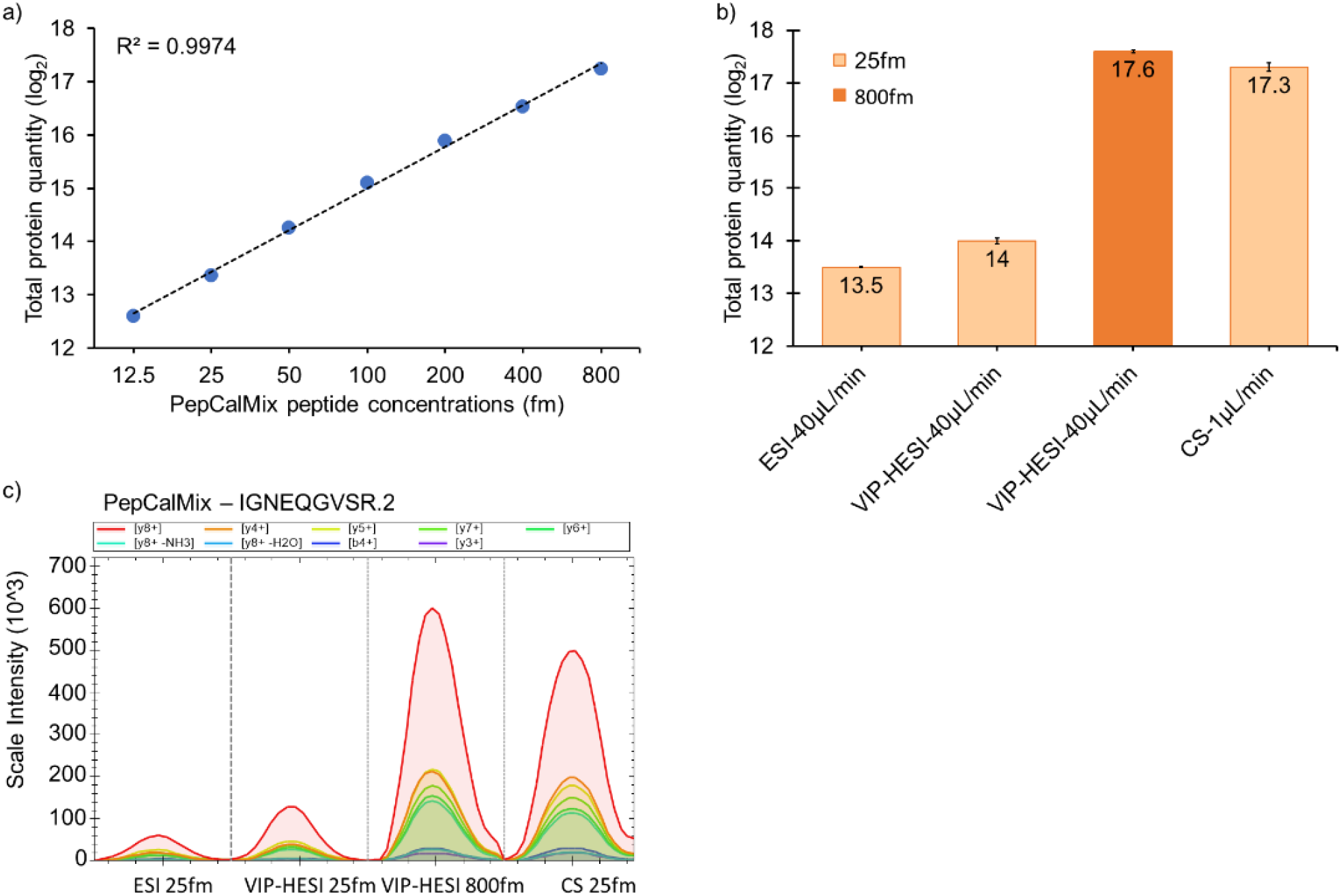
Evaluation of the linearity and sensitivity comparison of VIP-HESI, ESI, and CS ion source setups using PepCalMix measurements. **a)** Line graph of PepCalMix peptides to assess the linearity measured by dia-PASEF mode in timsTOF mass spectrometer. **b)** Bar plots of total quantities of all 20 PepCalMix peptides observed with ESI, CS ion sources at 25 fmol injection amount at 40 μL /min and 1 μL /min respectively, and with VIP-HESI source at 25 & 800 fmol injection amounts at 40 μL /min. Keeping the concentration of the eluting peak height constant (25 fmol for 1 μL = CS, 800 fmol for 40 μL for VIP-HESI), the response of VIP-HESI is even increased by 30% and a slightly higher peak response using the VIP-HESI source is observed. The error bars indicate the variability within three replicates represented as the standard error of the mean at the log2 scale. These are calculated as the ratio of the standard deviation of the peptide intensities observed with each source setup replicate to the square root of the sample size (*n* = 3). **c)** Extracted Ion Chromatogram (XIC) plots of a single precursor IGNEQGVSR.2 representing each source setup, colored by MS2 fragment intensity observed per measurement, as per the injection amount and flow rates.

Furthermore, to evaluate the comparative performance gains of VIP-HESI over the standard ESI and CS ion sources, we measured PepCalMix peptides at different dilution amounts. First, we compared the total protein quantity observed with the ESI to VIP-HESI sources using 25 fmol injection amounts at 40 μL/min flow rates. The VIP-HESI yielded 50% (log2=0.5) more total protein abundance than the standard ESI ion source as shown by higher peak areas with the same concentration and flow rate **(Fig. 1b, 1c, and Supplementary Table 4b))**. Subsequently, we acquired a 25 fmol PepCalMix amount using a CS source and compared it with a thirty-two-fold (800 fmol) higher injection amount measured using a VIP-HESI source to compensate for the loss of signal due to sample dilution at higher flow rates. The VIP-HESI source provided 30% (log2=0.3) more total protein quantity than the CS ion source, signifying better performance is achieved with the former source **(Fig. 1b and Supplementary Table 4 b))**. This is further evident by calculating the sensitivity factor (fmol/nL) by estimating the ratio of injected concentration to the flow rate for both sources. PepCalMix measurements of 800 fmol at 40 μL/min using VIP-HESI source provided a ratio of 0.02 fmol/nL whereas CS setup gave a ratio of 0.025 fmol/nL, indicating the former source is capable of achieving at least similar or higher sensitivity with higher injection amounts than its CS counterpart **(Fig. 1b)**. The effect of different injected amounts of PepCalMix peptides using different ion sources on the signal intensity of extracted MS2 chromatograms is exemplified with precursor IGNEQGVSR.2 **(Fig. 1c)**. These observations lead us to conclude that enhanced sensitivity is achieved with the VIP-HESI source as the result of its probe gas-assisted thermal desolvation which improves the spray performance under microflow rate conditions. This experimental setup is extremely beneficial for biological applications where the protein sample amounts are in abundance.

### Comparative performance of the VIP-HESI and CS ion sources using undepleted mouse plasma sample by dia-PASEF analysis

To demonstrate the comparative performance of both electrospray ion source units on more challenging samples where the total sample amount was not limited, we measured two biological replicates (with a single technical injection each) for a control UMHET mouse plasma sample with six different injection amounts and two different flow rates (see Experimental Procedures). Precisely, we studied the effects of sample amount of 0.4 μg at 1 μL/min using a 15 cm × 0.15 mm × 1.5 μm column coupled with CS and compared with 2 μg, 4 μg, 10 μg, 20 μg, and 40 μg sample amounts at 40 μL/min using a 15 cm × 1 mm × 1.7 μm Kinetix column (Phenomenex) coupled with VIP-HESI source. For a fair comparison between the two electrospray types, the same length of the column was used, packed with similar C18 particle sizes, and the flow rates were optimized based on the internal diameter of these columns.

dia-PASEF analysis of replicates using Spectronaut resulted in the identification of 5,795 - 8,610 precursors and 493 - 802 unique proteins at <1% protein FDR **(Fig. 2a, 2b, and Supplementary Table 5)**. The VIP-HESI showed an increase in precursors and protein identifications with more sample amounts being injected, although the gain was less prominent beyond the 20 μg injection amount indicating a signal saturation was attained **(Fig. 2a, 2b)**. Interestingly, injecting fifty times (20 μg) more sample amounts (with 40 times more sample dilution) using VIP-HESI, outperforms CS 0.4 μg measurement by identifying 3% more precursors and 12% more proteins **(Fig. 2a, 2b, and Supplementary Table 5)**. Next, we evaluated the robustness and reproducibility of quantitative measurements by comparing the precursor abundances of two biological replicate injections from these sources as estimated by Spectronaut. As expected for a high flow rate chromatography in VIP-HESI 20 μg measurement, a total of 7,074 precursors were quantified by both biological replicates with a high positive correlation (R^2^ = 0.94). With CS 0.4 μg replicate measurements, 6,387 precursors were quantified with a positive correlation (R^2^ = 0.89), indicating more variation was observed with nanoflow rates, compared to VIP-HESI-microflow runs **(Fig. 2c, 2d)**. Subsequently, we estimated the number of data points measured across the elution profiles and base peak widths for precursors identified in both VIP-HESI 20 μg and CS 0.4 μg measurements that used 1 mm and 0.15 mm ID columns, respectively. Similar median base peak widths of 0.16 and 0.14 minutes were observed with VIP-HESI 20 μg and CS 0.4 μg measurements, respectively. For both electrospray units, a median value of 5 data points per elution peak was obtained that provides an optimal quantification of an identified precursor using Spectronaut analysis (**Fig. 2e, 2f**). Similar results were obtained by injecting one hundred times (40 μg) more sample amount for analysis coupled with VIP-HESI **(Fig. 2a, 2b, and Supplementary Figure 1)**. These results demonstrate that the VIP-HESI source, combined with a 1mm micro-bore column using μLC–MS/MS, can be applied to complex tryptic digests that are not sample-limited, such as blood plasma, to achieve higher sensitivity and higher quantification precision of precursors and proteins.

**Figure 2.**
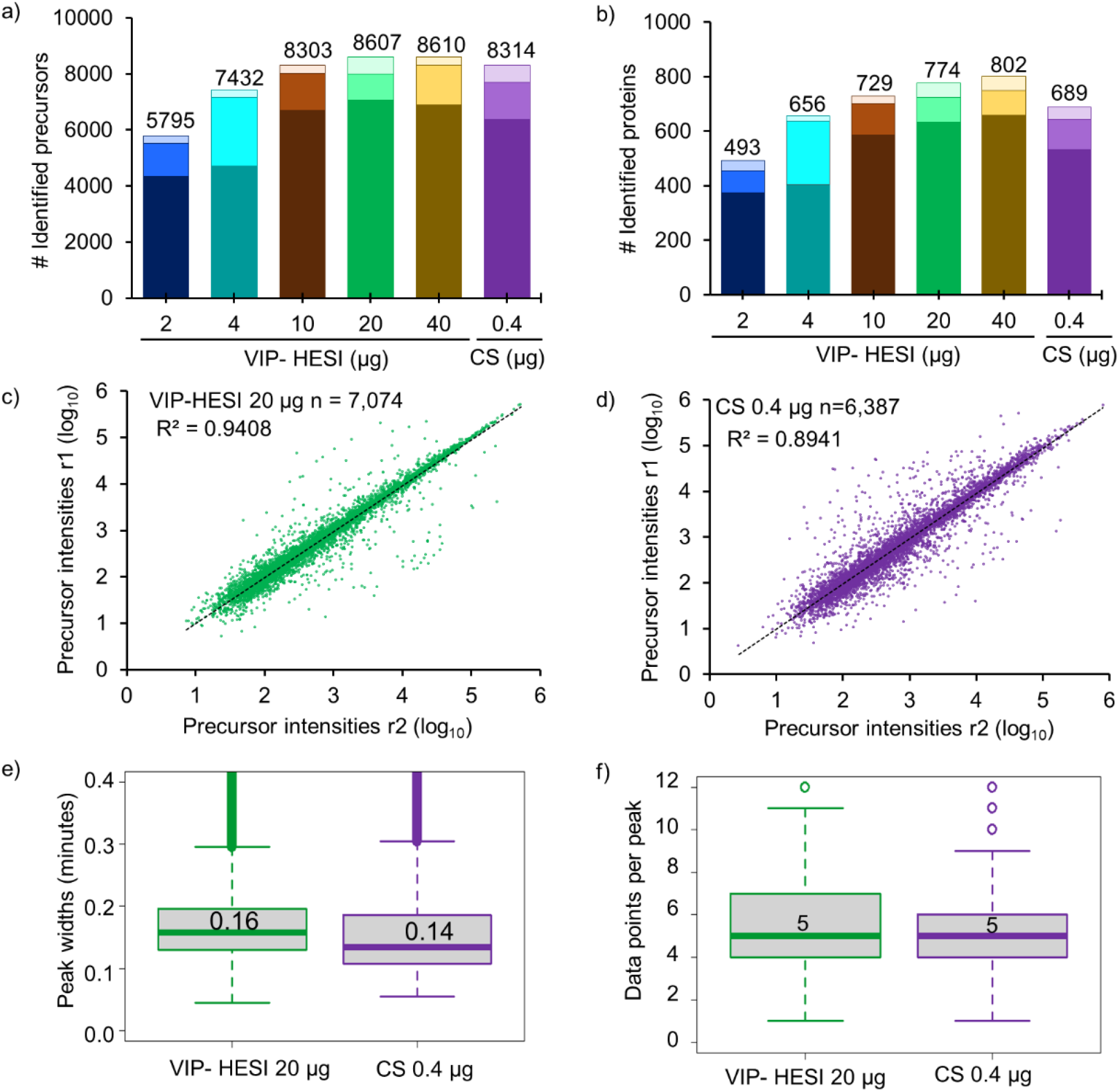
Comparative performance assessment of VIP-HESI and CS ion source setups using different injection amounts of undepleted mouse plasma samples. **a)** Number of precursors and **b)** protein groups identified with different injection amounts of mouse plasma samples analyzed in duplicate with a 45-minute 40μl/min microflow gradient. The darker color bar represents common identification between the replicates whereas the two lighter color bars at the top represents exclusive precursors and proteins identified in each biological replicate at the different concentration measurements. Pearson correlation of precursor intensity values obtained from 7,074 and 6,387 precursors that were quantified in both replicates (r1 and r2) using **c)** VIP-HESI and **d)** CS source setups respectively. **e)** Distribution of the base peak widths in minutes of precursors identified for VIP-HESI 20 μg and CS 0.4 μg measurements estimated by Spectronaut. A median value of 0.16 (9.6s) and 0.14 (8.4s) peak widths were observed for VIP-HESI 20 μg and CS 0.4 μg injections, respectively. The first and third quartiles are marked by a box with a whisker marking a minimum/maximum value ranging to 0.6 interquartile and the median is depicted as a solid line. **f)** Distribution of data points per elution peak for the VIP-HESI 20 μg and CS 0.4 μg measurements estimated by Spectronaut. The first and third quartiles are marked by a box with a whisker marking a minimum/maximum value ranging to 3 interquartile and the median is depicted as a solid line.

### Robust and quantitative analysis of hundreds of plasma proteomes by VIP-HESI source coupled with micro-flow liquid chromatography

To demonstrate the reproducible high-throughput application of VIP-HESI source, we analyzed 284 undepleted mouse plasma samples. Two consecutive blanks after each sample were injected for column washing purposes, and one quality control (QC) sample, K562 digested peptides, was acquired after every thirty-two plasma samples to assess the technical variation in instrument performance. In total there were 891 consecutive injections over the course of ~20 days in this experimental setup. Once established that the 20 μg of mouse plasma sample is the optimal amount for this setup from our earlier experiments (**Fig. 2**), we injected this amount for each sample using a 45-minute (3-35% ACN) HPLC gradient.

Large-scale dia-PASEF analysis with Spectronaut resulted in the identification of 17,628 precursors and 1,267 mouse plasma protein groups at <1% global protein FDR **(Fig. 3a, and Supplementary Table 6)**. Overall, we observed similar identification of precursors (9,328-11,885) and proteins (850-1,067) across all the samples, signifying the stable performance of the micro-flow LC system **(Fig. 3a)**. The retention times (RT) of the spiked in iRT peptides in 284 samples showed very high chromatographic reproducibility with median CV of 0.4% **(Fig. 3b and 3c)**. In addition, we checked the stability of the instrument setup by estimating the variation in the nine K562 QC samples. With 2 μg of K562 sample injection using a 21 minute gradient (3-35% ACN) identified an average of 4541 ± 5 protein groups based on 50,657 ± 437 precursors, signifying robust qualitative performance observed across QC samples **(Supplementary Figure 2 and Supplementary Table 7)**. Subsequently, we evaluated the quantitative precision between these QC technical replicates by estimating the coefficient of variation (CV) for the obtained protein quantities. The median CVs of the proteins quantified in nine technical replicates were below 10%, indicating an excellent reproducibility in quantitation across the whole experiment (891 injections) **(Supplementary Figure 2 and Supplementary Table 7)**.

**Figure 3.**
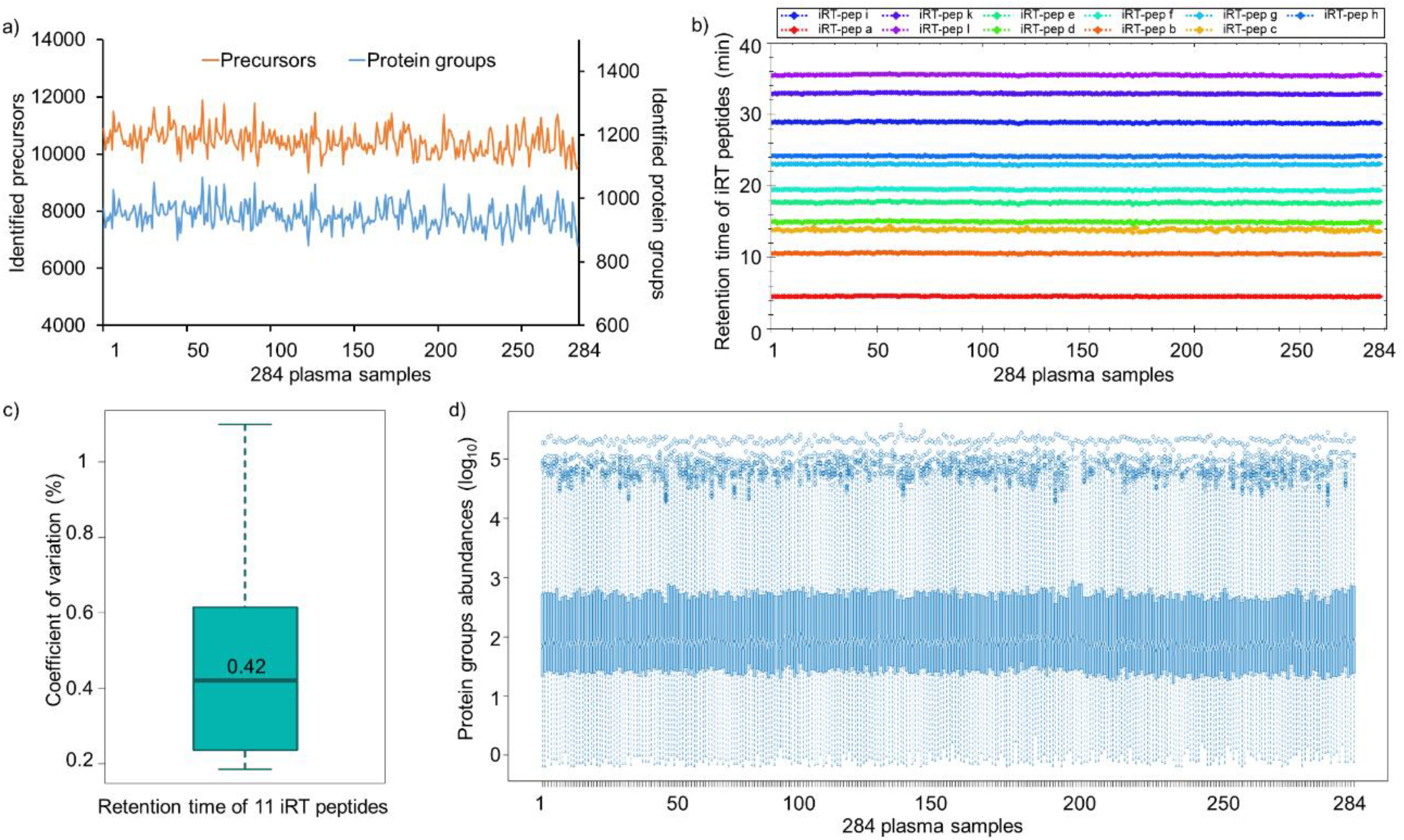
Large-scale quantitative analysis of undepleted mouse plasma samples using VIP-HESI coupled with microflow LC–MS/MS system. **a)** Number of precursors (orange) and protein groups (blue) quantified with 284 mouse plasma samples analyzed in a 45-minute 40μl/min microflow gradient on Vanquish Neo HPLC coupled to Bruker timsTOF Pro. **b)** A retention time QC plot for 11 iRT peptides spiked in each sample. **c)** Distribution of coefficient of variation (CV) in retention time of iRT peptides across 284 samples and the median is depicted as a solid line. **d)** Distribution plot of raw (un-normalized) protein quantities estimated in dia-PASEF analysis for 284 undepleted mouse plasma samples. Log10 protein quantities are plotted on the y-axis, sample number on the x-axis. Overall, protein quantities were similar across all samples with good alignment of sample medians and distributions over the first and third quartiles.

Plasma proteomics is challenging due to the high dynamic range of protein concentrations that could lead to the identification of mostly highly abundant proteins and poor detection for low abundant proteins resulting in less proteome coverage of these samples^27,28^. Further, we explored the variation of raw protein abundances across 284 mouse plasma samples and demonstrated that the distribution of protein quantities was similar across all samples even before normalization **(Fig. 3d)**. The medians were well-aligned across all the samples, indicating most proteins contributed equally to the observed total quantity in any sample, signifying consistency in the data acquisition with good ionization spray stability. As expected, protein quantities showed a large dynamic range across five orders of magnitude **(Fig. 3c)**. These results confirm the superior performance of VIP-HESI source with microflow chromatography for high-throughput applications that require reproducible results over extended periods of time.

### Performance of VIP-HESI microflow setup in a Slice-PASEF mode for analysis of low sample amounts

Lastly, we assessed the performance of microflow setup coupled with VIP-HESI source with the recently introduced Slice-PASEF, a continuous fragmentation mode of all precursor slices using trapped ion mobility^18^ and compared it to a standard dia-PASEF mode. Specifically, we evaluated its performance on the dilution series of a tryptic digest standard produced from the HeLa cell line, acquired in triplicates. We separated HeLa sample amounts of 10 ng, 100 ng, and 1000 ng at 20 μL/min flow rate with a 20 cm x 0.5 mm ID, ReproSil Pur 120 C18-AQ, *d*_P_ 1.9 μm (Dr. Maisch, GmbH) column setup and operated with a 21 minute chromatographic gradient (3-35% ACN). For contrast, we used the 1F Slice-PASEF scheme and compared it to a preformed dia-PASEF scheme featuring 25 Da isolation windows covering 400 to 1000 *m/z* range (see Experimental Procedures).

Slice-PASEF and dia-PASEF analysis with DIA-NN with different HeLa sample amounts resulted in the identification of 6,592 - 49,130 precursors at <1% precursor FDR and 1,769 - 6,343 protein groups at <1% protein FDR for the range of 10-1000 ng sample amounts. The Slice-PASEF with VIP-HESI microflow shows a significant increase in the number of precursor and protein groups, especially with low protein amount measurements **(Fig. 4a, 4b, and Supplementary Table 8)**. For 10 ng sample amounts, Slice-PASEF yielded 63% more precursors and 38% more protein groups than dia-PASEF. This is the result of a high MS/MS duty cycle in the 1F Slice-PASEF acquisition method, enhancing the sensitivity of the measurement. In contrast, with the higher injection amounts of 1000 ng, more ions are trapped and released by the TIMS device, the dia-PASEF method with a lower duty cycle gains more identifications than its counterpart as it optimally balances between the time spent on accumulating the spectra and selecting new spectra **(Fig. 4a, 4b, and Supplementary Table 8)**.

**Figure 4.**
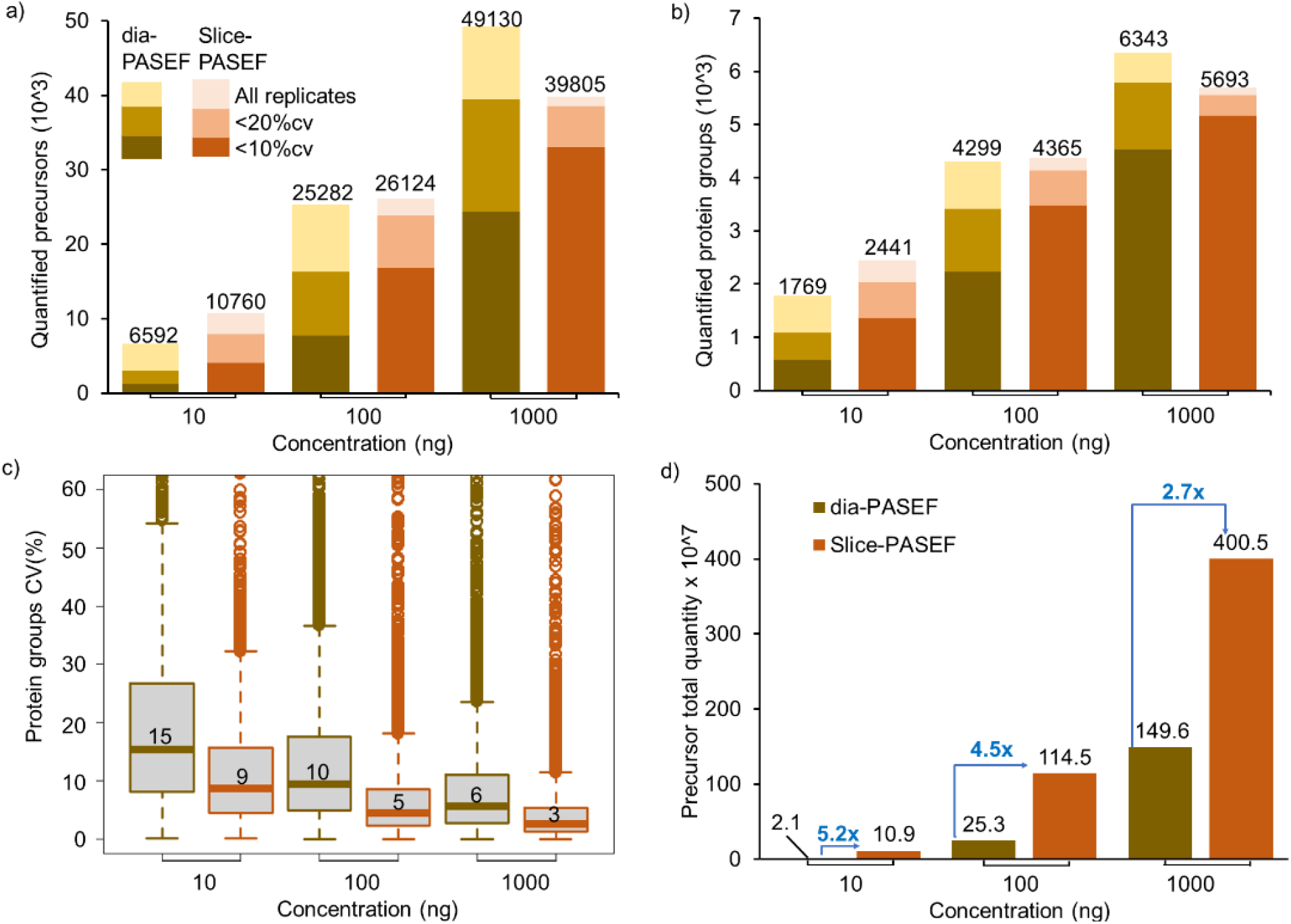
Performance of VIP-HESI-microflow setup with low sample amounts measured using dia-PASEF and Slice-PASEF acquisition methods. **a)** The number of precursors and **b)** protein groups quantified with different injection amounts of a HeLa tryptic digest analyzed in triplicates with a 21-minute 20μL/min microflow gradient on Vanquish Neo HPLC coupled to Bruker timsTOF Pro. The precursors and protein groups that were observed in all three replicates of each measurement were represented as all replicates and quantification precision expressed as coefficient of variation (CV) are illustrated at 10% and 20% levels. The Slice-PASEF method outperformed the dia-PASEF method in a 10ng sample amount by quantifying more precursors and proteins at both 10% and 20% CV levels. **c)** Distribution of the coefficient of variation (CV) of protein groups identified in all three replicates of different measurements at 1% protein FDR estimated by DIA-NN. The first and third quartiles are marked by a box with a whisker marking a minimum/maximum value ranging to 18 interquartile and the median is depicted as a solid line. **d)** The total quantity of all precursors identified in each measurement estimated by DIA-NN. The signal boost in the total quantity of all precursors between the same injection amounts is marked by the blue-colored arrows. The maximum boost is observed in the lowest sample amount of 10ng measurements using the Slice-PASEF method.

Further, at all injection amount measurements, the 1F Slice-PASEF method demonstrated more precise quantitation. This effect is more profound at low sample amount measurements of 10 ng where the Slice-PASEF method precisely quantified 2.4 times more protein groups with a coefficient of variation (CV) < 10% and 1.8 times more protein groups with a CV <20% than dia-PASEF acquisition **(Fig. 4b)**. Additionally, we evaluated the quantitative precision among different sample amount measurements using Slice-PASEF and dia-PASEF acquisition methods. The median CVs of the proteins quantified in three technical replicates of Slice-PASEF measurements were 8.7%, 4.6%, and 2.7% for 10 ng, 100 ng, and 1000 ng respectively. Whereas dia-PASEF quantitative variation among the replicates was more as median CVs were 15.3%, 9.7%, and 5.7% for 10 ng, 100 ng, and 1000 ng respectively **(Fig. 4c)**. On comparing the total precursor quantity estimated by DIA-NN for both acquisition methods, Slice-PASEF provided a significant signal boost from 5.2 fold increase for 10 ng to 2.7 fold increase for 1000 ng concentration than its counterpart **(Fig. 4d)**. These results indicate that an optimized VIP-HESI source in conjunction with microflow chromatography is capable of measuring low sample amounts with superior quantitation using Slice-PASEF mode in timsTOF MS.

## Conclusion

Here, we demonstrate the performance boost of the VIP-HESI source coupled with microflow chromatography flow rates and compared it to ESI and CS ion source units using a Bruker timsTOF MS. With the enhanced sensitivity, high ion-spray stability, and robust quantitative performance characteristics, VIP-HESI source is an efficient alternative to the nanoflow liquid chromatography approach for a wide range of proteomic applications.

## Data Availability

All mass spectrometry dia-PASEF raw folders corresponding to PepCalMix, K562, and HeLa experiments (.d), dda-PASEF raw folders of HeLa fractionated samples (.d), FASTA database used to generate the assay libraries (.fasta), spectral assay libraries (.txt), its DIALib-QC reports (.tsv) and output information from Spectronaut and DIA-NN as supplementary tables (.xls) have been deposited with the ProteomeXchange Consortium *via* the PRIDE partner^29,30^ repository with the dataset identifier PXD040116 (http://www.ebi.ac.uk/pride).

Username: Password: 0rEydxQk

## Supporting Information

**Supplementary Table 1**: Metadata of the mouse liver, kidney, gastrocnemius muscle tissues, and plasma samples.

**Supplementary Table 2**: Acquisition and LC separation details used in this study.

**Supplementary Table 3**: DIALib-QC report of spectral assay libraries “HeLa_timsTOF_DDA_DIA_Library” and “Mouse_tissue_plasma_DDA_DIA_Library”.

**Supplementary Table 4: a)** Linearity assessment of timsTOF using VIP-HESI source with different concentrations of PepCalMix peptidesTotal protein quantity. **b)** Comparative performance of VIP-HESI with standard ESI and CS ion sources using 20 synthetic PepCalMix peptides.

**Supplementary Table 5: a)** VIP-HESI Mouse plasma measurements. **b)** CS Mouse plasma measurements.

**Supplementary Table 6: a)** Mouse Plasma Precursor matrix. **b)** Mouse Plasma Protein matrix. **a)** Mouse Plasma RT-Chart Data.

**Supplementary Table 7: a)** K562 QC sample measurements analyzed by Spectronaut. **b)** Protein groups and **c)** Precursor sample matrices.

**Supplementary Table 8**: Comparative performance of VIP-HESI microflow setup using Slice-PASEF and dia-PASEF modes. **a)** dia-PASEF-10ng, **b)** dia-PASEF-100ng, **c)** dia-PASEF-1000ng, **d)** Slice-PASEF-10ng, **e)** Slice-PASEF-100ng, **f)** Slice-PASEF-1000ng.

## Contributions

R.L.M. conceived and designed the study. M.K.M. designed and performed dia-PASEF, Slice-PASEF MS data acquisition, spectral library generation, and data extractions using Spectronaut and DIA-NN tools. M.K.M prepared K562 samples. S.M. prepared HeLa samples. M.M. and D.B. prepared mouse tissue and plasma samples. M.K.M and T.J.P prepared PepCalMix dilutions. T.J.P. installed the CS, ESI, and VIP-HESI ion sources. M.K.M., C.K., and R.L.M. performed data analysis. M.K.M., C.K., and R.L.M. wrote the manuscript with contributions from all authors.

## Acknowledgments

This work was funded in part by the National Institutes of Health grants from the National Heart, Lung, and Blood Institute R01HL133135, the Office of the Director S10OD026936, the National Institute on Aging U19AG023122, and the National Science Foundation award 1920268. We thank Dr. Gary Kruppa (Bruker) and the technical resource team at Bruker for access to the VIP-HESI source used in this work. We thank Dr. Stephanie Kaspar-Schoenefeld (Bruker) for collecting MS data of fractionated HeLa lysate tryptic digest for spectral library generation.

## Ethics declarations

The authors declare no competing interests.

## TOC figure

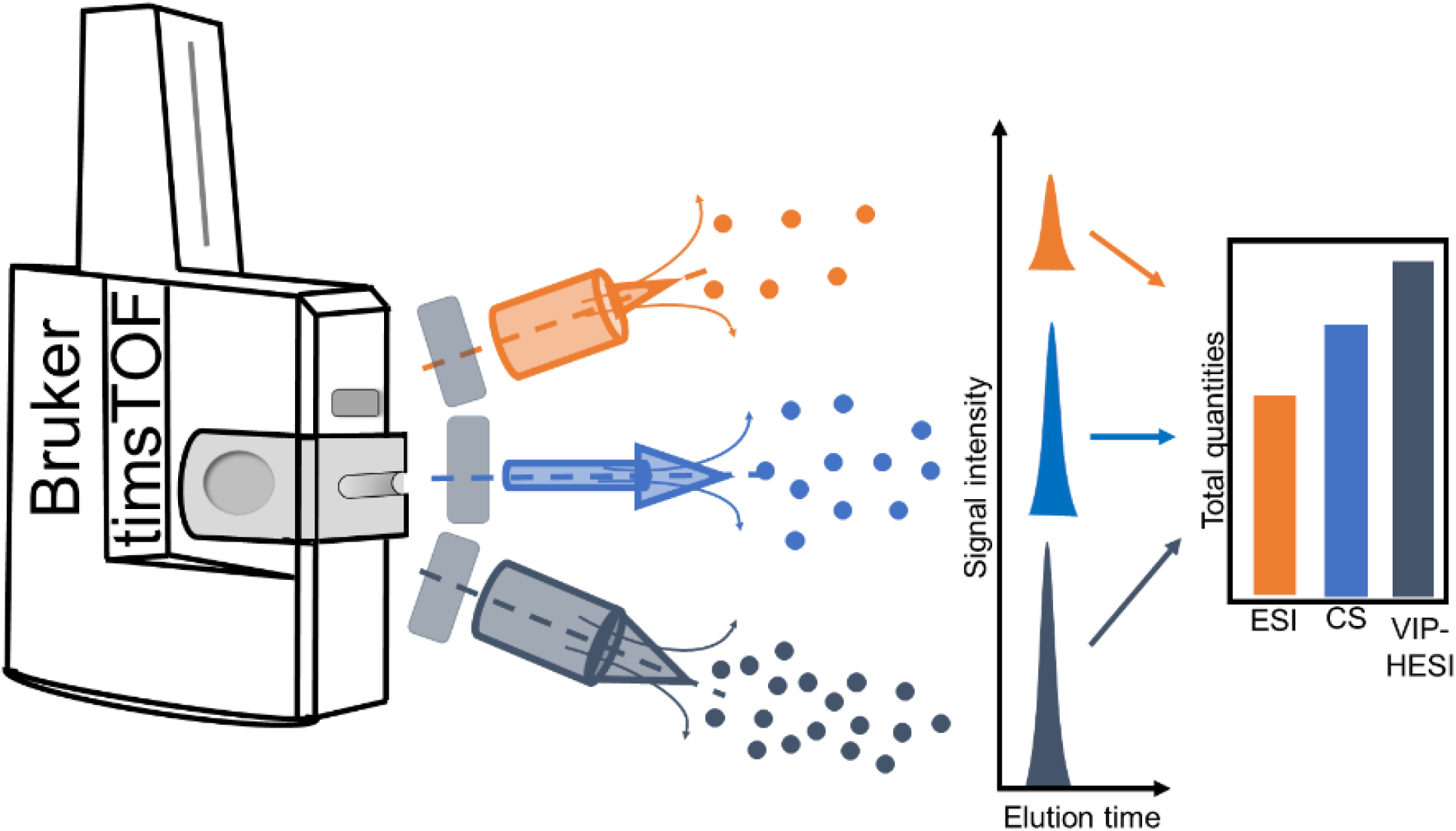

---

**Supplementary Figure 1.**
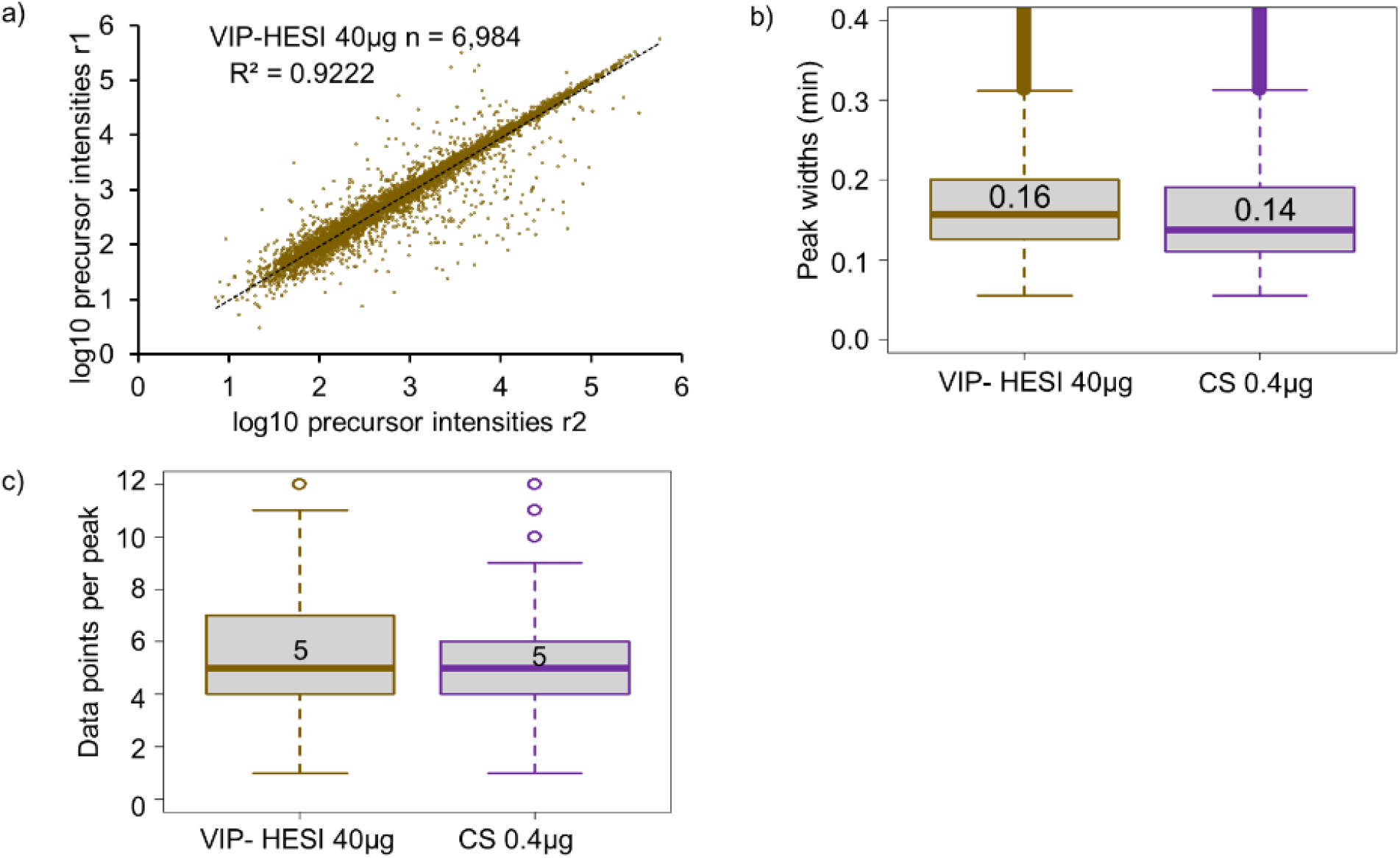
**a)** Pearson correlation of precursor intensity values obtained from 6,984 precursors that were quantified in both replicates using VIP-HESI 0.4 μg measurements of mouse plasma samples. **b)** Distribution of the base peak widths of precursors identified for VIP-HESI 40 μg and CS 0.4 μg measurements estimated by Spectronaut. A median value of 0.16 (9.6s) and 0.14 (8.4s) peak widths were observed for VIP-HESI 40 μg and CS 0.4 μg injections, respectively. **c)** Distribution of data points per elution peak for the VIP-HESI 40 μg and CS 0.4 μg measurements estimated by Spectronaut.

**Supplementary Figure 2.**
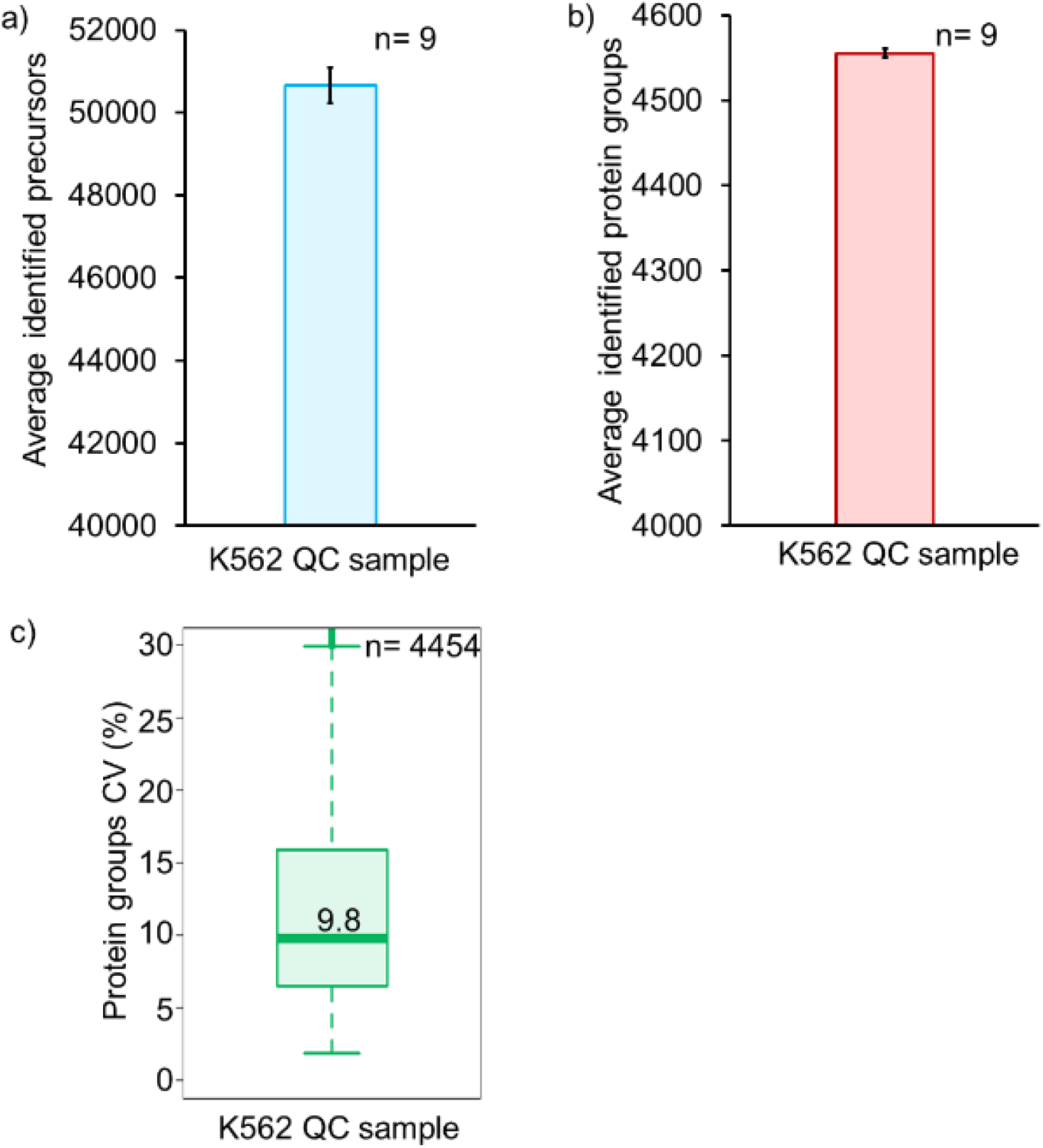
Estimation of MS instrument variation across the 891 VIP-HESI injections using K562 QC samples. The average **a)** number of precursors and **b)** protein groups identified with nine technical replicates. **c)** Distribution of coefficient of variation (CV) proteins identified in all nine replicates at 1% protein FDR estimated by Spectronaut. The median CV of 10% correlates well with the signal stability achieved using the VIP-HESI source setup. The first and third quartile are marked by a box with a whisker marking a minimum/maximum value ranging to 10 interquartile and the median is depicted as a solid line

